# *IsoNet2* determines cellular structures at submolecular resolution without averaging

**DOI:** 10.64898/2025.12.09.693325

**Authors:** Yun-Tao Liu, Hongcheng Fan, Jonathan Jih, Liam Tran, Xiaoying Zhang, Z. Hong Zhou

## Abstract

We introduce *IsoNet2*, an end-to-end self-supervised deep-learning method that directly reconstructs high-quality 3D densities from cryogenic electron tomography. A unified network simultaneously performs denoising, contrast transfer function correction, and missing-wedge restoration, achieving ∼20 Å resolution without averaging. A feature-rich GUI enables rapid, dataset-specific fine-tuning for end-users. *IsoNet2* resolves domain organization in HIV capsid proteins, tRNA occupancy in individual ribosomes, and *in situ* architectures of mitochondrial respiration-related complexes, enabling atomic-level interpretation of cellular environments.

## Main

Two well-established averaging strategies—single-particle analysis (SPA) in cryogenic electron microscopy (cryoEM) and subtomogram averaging (STA) in cryogenic electron tomography (cryoET)— have transformed structural biology by enabling near-atomic-resolution reconstructions of macromolecular complexes. However, both approaches rely on a strong “single-particle” prior: the assumption that molecules being averaged are structurally identical or share a rigid core. This assumption is critical for achieving three objectives central to biological EM structure determination: (i) merging information from differently oriented particles to complete Fourier sampling, (ii) reducing noise through averaging, and (iii) enabling accurate contrast transfer function (CTF) correction. Yet many biologically important systems do not exist in large identical copies in cells. Pleomorphic assemblies, most nucleic acids, lipids, intrinsically disordered regions, phase-separated condensates, and mesoscale molecular organizations therefore remain “averaging-invisible,” largely inaccessible to the current paradigm of high-resolution structure characterization.

Efforts to improve interpretability of raw cryoET tomograms without averaging have included iterative reconstruction algorithms such as SIRT^1^ and SART^2^, which enhance contrast but sacrifice high-frequency detail. Early deep-learning approaches such as *CryoCARE*^3^ introduced *Noise2Noise*^4^ denoising, and the original *IsoNet*^5^ demonstrated that neural networks, when trained on user data, can produce effective missing-wedge compensation. Recent methods, including *DeepDeWedge*^6^ and *CryoLithe*^7^, integrate *Noise2Noise* training with wedge-aware design, but have yet to optimize for recovery of high-resolution 3D information, a necessary precursor to “true” single-particle analysis.

We now present *IsoNet2*, which resolves variation-faithful submolecular features in an authentic sample context, to move molecular structural interpretation beyond the traditional SPA/STA paradigm by eliminating obligate need for particle averaging. To accomplish this, we designed *IsoNet2* to integrate the three core objectives of cryoEM reconstruction in a single deep-learning optimization loop (Fig.1a and Extended Data Fig.1), using a “Petronas” architecture (two prongs communicating via bridge, reminiscent of Southeast Asia’s Petronas Towers) that implements (i) accurate missing-wedge compensation to restore Fourier completeness, (ii) *Noise2Noise*-driven denoising, and (iii) network-based CTF correction that sidesteps limitations of classical Wiener deconvolution, thereby facilitating recovery of high-spatial frequencies. Our method operates direct on even–odd tomograms and is implemented via a rich web- technology–based graphical user interface (Fig.1a and Extended Data Fig.2), replete with job submission management, live process monitoring, and user-definable network models and parameters. This enables robust and efficient dataset-specific tuning—an increasingly recognized approach to optimize model performance on small data regimes^8,9^ (as is often the case with unique or rare events visualized in cryoET).

**Figure 1.**
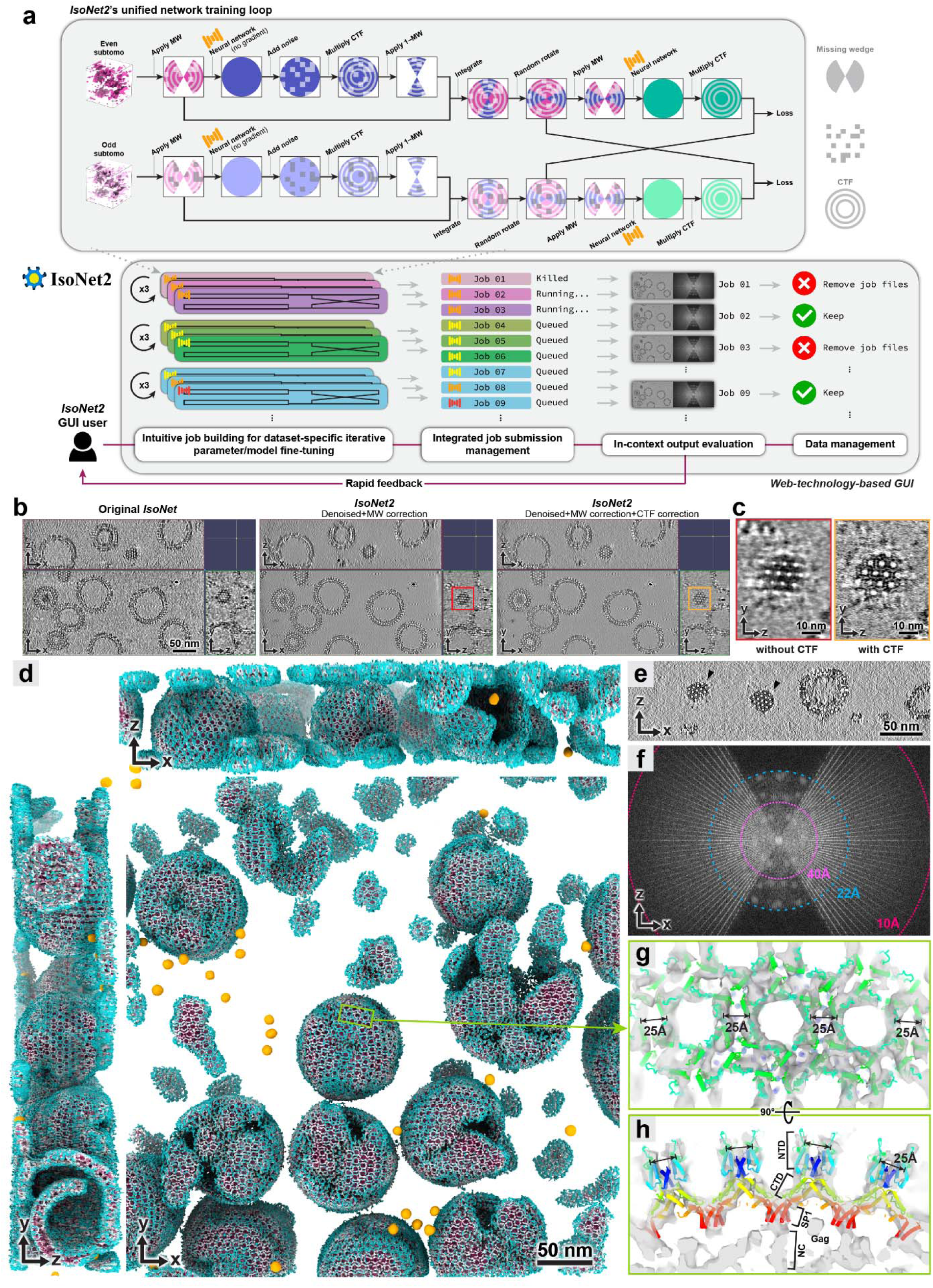
*IsoNet2*’s “Petronas” network architecture and full-featured GUI permit recovery of submolecular features **a,** *IsoNet2*’s bridged dual-prong training scheme operates in a self-supervised, end-to-end loop. A user-friendly GUI facilitates efficient dataset-specific fine-tuning and management of preprocessing and production runs. **b,** Tomographic slices of immature HIV particles (EMPIAR-10164)^15^ processed by *IsoNet1* and *IsoNet2* without/with CTF correction. **c,** Y-Z capsid slices boxed in (**b**) show clear lattice spacing in CTF-corrected. **d,** Orthogonal 3D renderings of *IsoNet2*-processed tomogram (CTF-corrected). Capsid densities colored by distance to lattice surface. Gold spheres are fiducial markers. **e-f,** X-Z slice (**e**) and its Fourier transform (**f**) show peaks (white spots) corresponding to lattice (black arrowheads) recovered in missing-wedge out to ∼22 Å. **g-h,** Top (**g**) and cross-section (**h**) views of HIV Gag atomic models (rainbow pipe-and-plank) fit in tomogram density. NTD: N-terminal domain; CTD: C-terminal domain; SP1: spacer peptide 1; NC: nucleocapsid.

**Figure 2.**
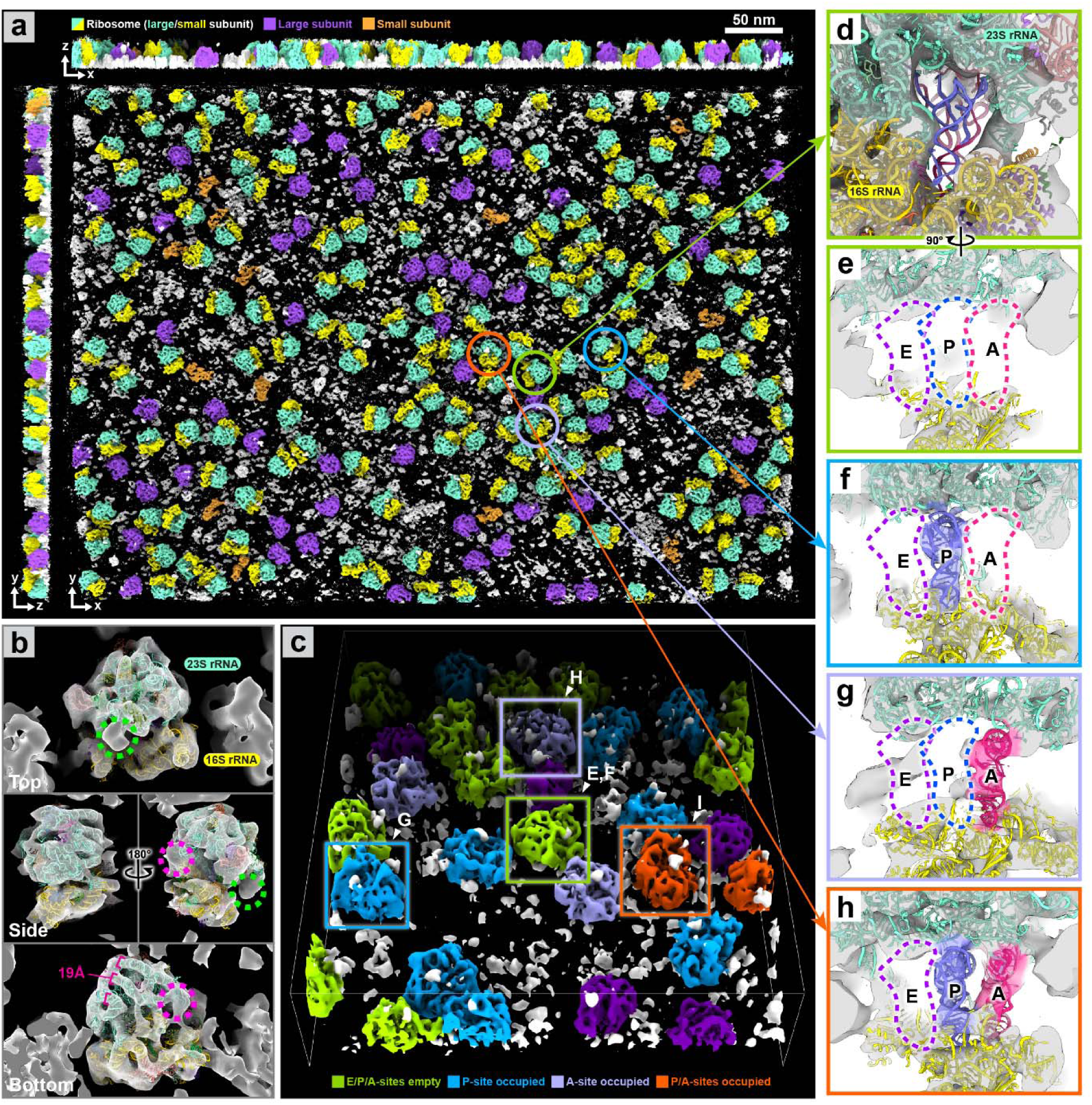
Direct resolution of ribosomal tRNA occupancy by *IsoNet2* **a,** *IsoNet2*-processed tomogram of purified 70S ribosomes (EMPIAR-10985)^17^, colored by ribosomal subunit. **b,** Top, side, and bottom views of unaveraged ribosome docked with PDB-7K00. Dashed circles indicate resolved rRNA density invisible in averaged reconstructions. **c,** Visual classification of ribosomes by tRNA occupancy: empty (green), P-site occupied (blue), A-site occupied (lavender), P/A-site occupied (orange). **d–h**, Observed tRNA binding states in E/P/A sites. In (**d**), E/P/A sites are empty, but tRNA models shown for reference.

For missing-wedge correction, *IsoNet2* deploys a self-supervised strategy where subtomograms first are randomly extracted, then passed through a neural network in inference mode to produce initial volume predictions. Initial predictions are filtered with corresponding CTF and noise, after which the missing-wedge region of these filtered volumes is integrated with original data to generate missing- wedge–“filled” subtomograms. These “filled” subtomograms contain experimental information in the measured Fourier region and network-predicted information in the missing region. “Filled” subtomograms are randomly rotated and used as training targets, where their missing-wedge–applied versions serve as inputs, allowing a U-Net–based architecture (Extended Data Figs.1c) to learn missing-wedge restoration across all orientations. Critically, unlike the original *IsoNet* [henceforth *IsoNet1*], which assumed a fixed ±60° tilt range, *IsoNet2* supports per-tomogram wedge geometries, making it suitable for FIB-milled lamellae with variable tilt ranges.

For denoising, *IsoNet2* adopts *Noise2Noise*^2^ techniques using statistically independent half-datasets derived from even–odd movie frames or tilt images. By minimizing discrepancies between the two halves, *IsoNet2* removes uncorrelated noise while preserving signal, similar to implementations in *CryoCARE*^3^, *WARP/M*^10^, and *Topaz-Denoise*^11^. To separate the loss between missing-wedge compensation and *Noise2Noise* denoising, we adapted a masked loss strategy first demonstrated in *DeepDeWedge*^6^.

CTF correction in *IsoNet2* is performed by exploiting the network’s predictive capacity. Namely, the network output is multiplied by experiment-specific CTF before comparison against intrinsically CTF-modulated targets (from “filled” tomograms). This forces the network to output pre**-**CTF–applied (i.e., CTF-corrected) predictions, avoiding traditional Wiener filtering while recovering information near CTF zeros. To address deviations of low-frequency signal from theoretical CTF behavior, we clamp the CTF curve to unity below the first peak, similar to the “CTF intact first peak” approach in *RELION*^12^. We also implement user-defined B-factor weighting, allowing tailored training to prioritize high- or low-resolution information recovery (Extended Data Fig.1b).

All components—missing-wedge restoration, denoising, and CTF correction—are jointly optimized as differentiable operations in a fully end-to-end training loop. This unified approach delivers substantially improved tomogram quality and reduces over-smoothing effects observed in *DeepDeWedge* reconstructions (Extended Data Fig.3). Unlike *IsoNet1*, *IsoNet2* eliminates iterative training–prediction– regeneration cycles, eliminates explicit particle extraction and external CTF deconvolution, and adopts mixed-precision training^13^, accelerating *IsoNet2* processing by roughly an order of magnitude (Extended Data Fig.1d). The resulting efficiency enables training with substantially larger subtomograms (typically 96^3^ to 128^3^ voxels versus 64^3^ in *IsoNet1*) and an additional down/upsampling layer in the U-Net^14^ architecture, providing a wider receptive field, all while permitting routine use of smaller pixel sizes (∼5 Å/pixel versus ∼10 Å/pixel in *IsoNet1*) for high resolution reconstructions.

**Figure 3.**
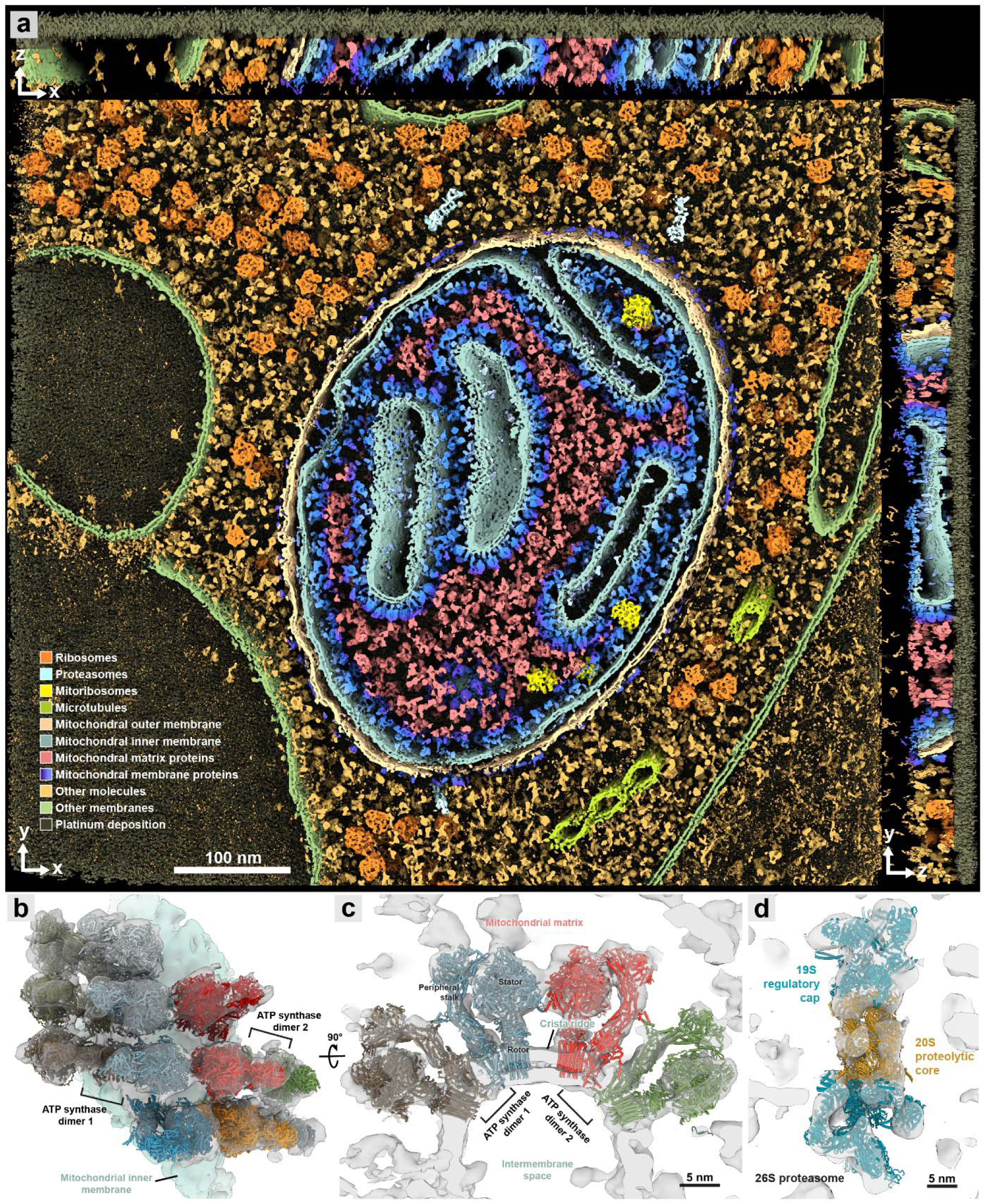
*IsoNet2* enables native visualization of molecular sociology at uniform threshold **a,** A “Goodsell-esque” 3D orthogonal rendering of an *IsoNet2*-processed FIB-milled tomogram of *C. reinhardtii* (EMPIAR-11830)^20^ shows the dense molecular environment in a cellular lamella. **b-c,** Top (**b**) and cross-section (**c**) view of ATP synthase dimers on a crista ridge. Atomic models of ATP synthase (individual dimers labeled “1” and “2”) are fit in unsegmented tomographic density without averaging. **d,** Tomographic density of a cytosolic 26S proteasome fit with atomic model.

We evaluated *IsoNet2* on an immature HIV virus-like particle dataset (EMPIAR-10164)^15^ previously used to benchmark *IsoNet1* (Fig.1b). These particles feature Gag proteins forming an incomplete spherical lattice of curved, hexagonally packed capsomers. *IsoNet2* dramatically enhanced tomogram clarity, revealing the narrow inter-hexamer gap created by capsid N-terminal domain (CA-NTD) ridges—a feature obscured without CTF correction (Fig.1c). Processed tomograms exhibited near-isotropic resolution, faithfully resolving the Gag lattice across orthogonal views, with lattice defects and local heterogeneity readily apparent (Fig.1d and Supplementary Video 1). In Fourier space, distinct lattice peaks extended to ∼22 Å even within the original missing-wedge region (Fig.1e–f), consistent with the ∼21 Å median local resolution assessed by *ResMap*^16^ (Extended Data Fig.4). Restored densities further delineated both CA-NTD and CA-CTD domains, the six-helix CA–SP1 bundle beneath each capsomer, and additional density comparable in strength to the capsomer directly beneath the CA–SP1 bundle, likely corresponding to the nucleocapsid domain for which no atomic model exists (Fig.1g–h).

We next applied *IsoNet2* to tomograms of purified 70S ribosomes acquired by *PACEtomo*^17^ (EMPIAR-10985; Extended Data Fig.1a). Processed volumes clearly resolved large and small ribosomal subunits (Fig.2a). Rigid-body fitting a 70S atomic model^18^ accurately placed double-stranded rRNA helices (∼20 Å diameter) in density (Fig.2b) in agreement with local resolution estimates (Extended Data Fig.5). Flexible peripheral rRNA extensions—typically lost during subtomogram averaging—were retained. Using *ChimeraX* virtual reality tools^19^, we directly inspected individual ribosomes in VR to classify them by native tRNA occupancy at the A, P, and E sites. *IsoNet2* yielded unambiguous tRNA densities, revealing a heterogeneous ribosomal population with A-site–only, P-site–only, A/P, or empty tRNA occupancy states (Fig.2c–h).

Finally, we applied *IsoNet2* to an *in situ* cryoET dataset of *Chlamydomonas reinhardtii* lamellae prepared by cryo-plasma FIB milling (EMPIAR-11830)^20^, analyzing fifteen mitochondria-containing tomograms (Extended Data Fig.6). Compared to *CryoCARE*, *IsoNet2* reduced missing-wedge artifacts and improved denoising (Extended Data Fig.7). Notably, *IsoNet2*’s ability to produce relatively uniform density quality across entire tomogram volumes permits visualization of all cellular components at a single threshold, creating a “Goodsell-esque”^21^ molecular panorama evocative of David Goodsell’s illustrations that vividly highlights authentic molecular crowding and sociology in cells (Fig. 3a and Supplementary Video 2). Macromolecular assemblies, including ribosomes, microtubules, and cristae- studded oxidative phosphorylation machinery, were instantly recognizable in 3D, permitting direct visualization of higher order architecture. For instance, ATP synthase, with stalk and stator regions clearly visible, presents as dimers arranged in semi-helical ribbons along crista ridges, consistent with prior STA^22^ (Fig. 3b–c and Extended Data Fig.8). At ridge apices, double rows of adjacent dimers are frequently observed, their spacing likely driving the characteristic 180° membrane curvature. Lastly, *IsoNet2* recovered sufficient detail to resolve three dispersed cytosolic 26S proteasomes^23^—distinguishing the 20S core, 19S regulatory caps, and subunit-level features (Fig.3d)—demonstrating that our network does not rely on copy number in sampled subtomograms to recapitulate accurate 3D structure.

Together, *IsoNet2*’s results demonstrate that a unified deep-learning strategy can achieve high-fidelity structural interpretation direct from raw tomographic data, without the need for particle averaging. Our streamlined GUI encourages robust end-user experimentation to extract optimal model performance tailored for individual data. When combined with predictive tools such as *AlphaFold*^24^ and immersive visualization frameworks in VR/AR, *IsoNet2* moves the field measurably closer to the long-standing goal of interpreting molecular sociology^25^ at near-atomic detail, to ultimately inform construction of comprehensive, full-atom models of cells.

## Methods

### Overview of *IsoNet2*

*IsoNet2* was implemented in Python and runs under the Linux operating system. The software package, including source code and documentation, is available on GitHub (https://github.com/IsoNet-cryoET/IsoNet2). *IsoNet2* uses the PyTorch^26^ deep-learning framework, replacing the TensorFlow^27^ backend previously used in *IsoNet1*.

*IsoNet2* unifies the three essential tasks of cryoEM tomogram reconstruction within a single optimization pipeline: robust missing-wedge compensation restoring Fourier completeness, *Noise2Noise*- based denoising, and a learned contrast transfer function (CTF)-correction module that circumvents Wiener deconvolution. *IsoNet2* also retains all core functionalities from *IsoNet1* for processing tomograms without even–odd splitting—including mask generation, CTF deconvolution, and the full ‘Refine’ workflow. In addition, *IsoNet2* introduces a *CryoCARE*^3^-like *Noise2Noise* denoising module for rapid denoising, which can optionally incorporate the network-based CTF-correction to yield higher-resolution outputs.

The software accepts paired tomograms (even and odd) as input, together with imaging parameters such as electron voltage, spherical aberration, amplitude contrast, defocus values, and tilt range. These parameters are required for CTF correction and missing-wedge correction. The STAR file format used in *IsoNet2* follows the *RELION-5*^28^ standard, ensuring compatibility with established cryoEM processing pipelines.

To generate paired tomograms, when the tilt images are acquired as dose-fractionated movies, even–odd splitting can be performed during motion correction: alternating movie frames (e.g., even vs. odd frames) are separated, motion-corrected, and summed independently, yielding two sets of images with same signal but statistically independent noise. If dose-fractionated movies are not available, even– odd tomograms can instead be generated directly from the tilt series by dividing the tilt images into two interleaved subsets (e.g., tilts 0, 2, 4… and 1, 3, 5…) and reconstructing each subset independently. This produces a pair of tomograms that share the same underlying structure but contain uncorrelated noise, which is essential for *Noise2Noise* learning in *IsoNet2*. We encourage users to perform frame-based even–odd splitting where possible.

### Graphic user interface (GUI)

*IsoNet2* includes a rich desktop graphical user interface (GUI) written in JavaScript, providing an integrated visual workflow for running and managing *IsoNet2* (Extended Data Fig.2). Built with Electron, React, and Node.js, the GUI combines a lightweight desktop front end with a Python backend executed within a Conda environment. It offers tools for dataset organization, parameter configuration, job submission, and real-time process monitoring, with streaming logs and progress visualization. The interface arranges the main processing steps in a left-hand menu, while the central panel shows the program’s live output during a run. Parameter drawers on the right allow users to select input files, adjust basic settings, and submit jobs. Asynchronous inter-process communication (IPC) between the Electron frontend and Python backend ensures responsive interaction even during GPU-intensive computation, supporting user-definable network models and parameter tuning.

### The neural network architecture of *IsoNet2*

*IsoNet2* employs a 3D U-Net architecture (Extended Data Fig.1c) similar to that of *IsoNet1* and *spIsoNet*^29^, and the basis of which is widely used in biomedical image restoration and segmentation^14^. Each convolutional block contains three 3D convolutional layers (kernel size 3×3×3) with leaky ReLU activations. The encoder path consists of four such blocks, each followed by a strided convolution layer (2×2×2) that halves the spatial dimensions while doubling the number of feature channels. The decoder path mirrors this structure, using transpose convolutions for up-sampling. Skip connections concatenate feature maps of equal resolution between the encoder and decoder paths to preserve high-resolution details. Compared with *IsoNet1*, the default network in *IsoNet2* increases network depth from three to four down-sampling levels while using a lighter 32-filter base.

By default, IsoNet2 uses mixed-precision technology13, providing up to a two-fold speed increase without loss of accuracy (Extended Data Fig.1d). The network is represented as in the following sessions, where denotes all trainable weights updated during training.

### End-to-end Refine process

End-to-end refinement is a defining and central process in *IsoNet2*, whereby input tomograms and imaging parameters inform training of a neural network that simultaneously performs denoising, missing-wedge compensation, and network-based CTF correction. Training typically uses around 3,000 subtomograms with cube sizes of 96^3^, and runs for approximately 50–100 epochs.

Unlike *IsoNet1*, which required separate iterative cycles for data updates and network refinement, *IsoNet2* integrates all operations within a single unified optimization loop. This streamlined design eliminates the need for intermediate manual steps such as subtomogram extraction and CTF deconvolution, vastly simplifying GUI integration and enabling the entire refinement workflow to be executable by a single command. For example:

*isonet.py refine tomograms.star --CTF_mode network*

### Refine step 1: Subtomogram preparation

Tomograms are first loaded and preprocessed as follows. The mean and standard deviation are computed from the central 64 slices of each tomogram. Subtomograms (default size 96³ voxels and at least 64³ voxels) are randomly cropped from even–odd tomograms. The number of subtomograms is user-defined and typically totals approximately 3000. Subtomogram extraction is incorporated into the training loop instead of as a separate function as in *IsoNet1*^5^ and *DeepDeWedge*^6^. Before feeding into the network, cropped subtomograms are normalized by subtracting the calculated mean and dividing the standard deviation values from the corresponding tomogram. When a mask is provided, the centers of the cropped subtomograms are restricted to masked areas. The CTF and optional Wiener filter volumes are also precomputed in this step, which will be described in the CTF section.

### Refine step 2: Generating missing-wedge–filled subtomograms

Self-supervised learning in *IsoNet2* relies on accurately generating the training targets that resemble “ground truth” from experimental data. In order to generate these targets, the extracted subtomograms first pass through an initialized neural network (which will be updated in the training loop) to generate predicted subtomograms: where even and odd subtomograms extracted at identical coordinates are denoted as and respectively, and is the index of the subtomogram.

These predicted subtomograms are subsequently added with noise and filtered with CTF that matches the noisy original subtomograms. We then generate missing-wedge–filled subtomograms by adding the information of the missing-wedge region of the predicted-and-filtered subtomograms to the original subtomograms, generating a set of subtomograms that are noisy and CTF modulated but missing-wedge partially filled. The formular of generating these subtomograms are as follows:

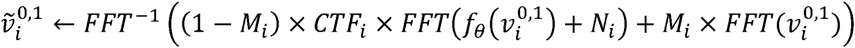

Where FFT stands for fast Fourier transform, is the missing-wedge mask which has value zeros in the wedge region and ones otherwise, is the noise volume that samples from normal distribution with standard deviation of : ∼ 0, , and Var means variation.

The neural network model parameters used in this step are not updated as no gradients are calculated. This step resembles the predict subtomograms step in *IsoNet1*’s refine loop, although *IsoNet1* does not have adding noise and CTF multiplication.

### Refine step 3: Rotating and applying missing-wedge on network input

Missing-wedge information is recovered by learning information from other orientations of the subtomograms. To generate rotated subtomograms and construct paired training samples with and without missing-wedge, we processed the missing-wedge–filled subtomograms using the formular as follows:

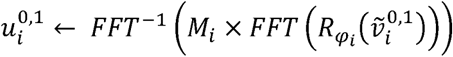

Where ! means to rotate a subtomograms with a randomly chosen 3D angle . , will be used as network input.

### Refine step 4: CTF correction

*IsoNet2* offers two options for CTF correction: 1. network-based and 2. Wiener filter based (*cf.* Fig.1a and Extended Data Fig.1f).

*CTF correction by the network*. After input subtomograms (containing the missing-wedge artifacts) are processed by the neural network, the output is then multiplied by CTF. The loss is computed between these CTF-multiplied outputs and the corresponding target rotated subtomograms. This forces the network to output pre**-**CTF–multiplied (i.e., CTF-corrected) predictions, avoiding traditional Wiener filtering while recovering information near CTF zeros.

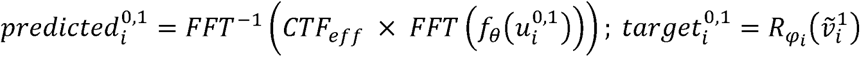

In the very low-resolution regime (lower than about 50 Å resolution) of the Fourier space of cryoET tomograms, the theoretical CTF does not faithfully reflect the actual CTF modulation present in the data^30^. To address this, we ignore the lowest-resolution region by generating a CTF curve only up to the first CTF peak (Extended Data Fig.1b), similar to the concept of the “CTF intact first peak” used in the *RELION*^12^ implementation. This modification avoids applying incorrect CTF amplitude corrections in the low-resolution range, which helps preserve the initial contrast of the original tomogram and leads to reconstructions with better high-resolution details.

The CTF curve is optionally accompanied by a user-defined B-factor parameter (Extended Data Fig.1b). This B-factor controls the frequency-dependent signal attenuation applied to the tomogram. A larger B-factor means the signal is attenuated faster with the increment of spatial frequency. We recommend setting the B-factor parameter to 200-300 for tomograms of isolated particles and B-factor to 0 for cellular tomograms. Considering the CTF intact first peak and B-factor, the effective CTF is as follows, where is the spatial frequency corresponding to CTF first peak:

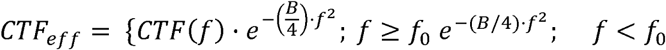

2. CTF correction through Wiener filter, with an empirical spatial signal-to-noise ratio (SSNR), as has been described in *IsoNet1*^5^ and *Warp*^31^:

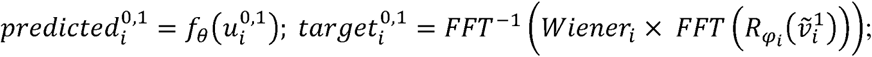

### Refine step 5: Compute loss

Network input subtomograms from both even and odd tomogram sets are normalized to match their original subtomograms and then passed through the network, and their corresponding outputs are used to compute the loss for training. To enable *Noise2Noise*-based denoising, predicted even subtomograms are compared against odd target subtomograms, and vice versa. The final loss is the average of these two comparisons:

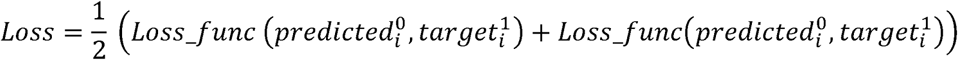

The loss function can be a standard L1 or L2 loss, or the masked loss introduced in *DeepDeWedge*. The masked loss is defined as:

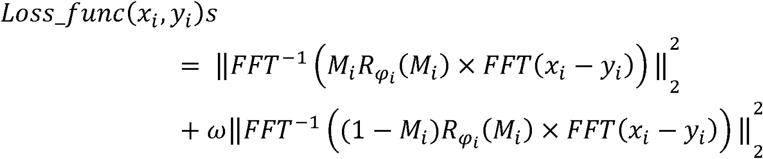

Where is the weighting factor that enables placing different weight on the missing-wedge correction and denoising. Larger means prioritizing the missing-wedge correction over *Noise2Noise* denoising. In practice, we found that using masked loss with high missing-wedge weight yields better performance in terms of the high-resolution feature preservation.

### Predict

After the Refine step, the trained network is saved as a model file with a file extension of “.pt”, which is then applied to the original tomograms or to other tomograms acquired under similar imaging conditions. The even and odd tomograms are predicted independently and combined to produce corrected tomograms

(Extended Data Fig.1e). Because processing full-size tomograms typically exceed GPU memory limits, the tomograms are divided into smaller 3D chunks. The network processes each chunk independently, and outputs are stitched together to generate the final corrected tomogram. In *IsoNet2*, the chunk size is set to match the subtomogram size (default is 96^3^) used during training.

To prevent visible boundaries between chunks—caused by limited context near their edges—we use an overlap-tile strategy^14^, where overlapping regions are jointly predicted and smoothly blended. By default, the overlap width is one quarter of the subtomogram. Because we implemented global normalization on tomograms (rather than normalizing individual subtomograms as in *IsoNet1*), patchy artifacts sometimes observed in *IsoNet1* reconstructions are eliminated. This prediction step is implemented in GUI or with a single command: *isonet.py predict tomograms.star network.pt*

### Other modules in *IsoNet2*

*IsoNet2* also includes a *Noise2Noise*-style denoising module similar to *CryoCARE*^3^. This denoising procedure is faster than the Refine step, making it useful for quick dataset assessment and for generating masks. In addition, *IsoNet2*’s denoising can optionally include CTF correction, producing higher-resolution denoised outputs. This denoise module can be performed in GUI or with the command: *isonet.py denoise tomograms.star*.

For tomograms lacking even–odd tomogram pairs, *IsoNet2* preserves the deconvolution-based methods and refinement workflow from *IsoNet1*. Therefore, denoising and missing-wedge correction for tomograms without even–odd pairs can be performed in a manner similar to the original *IsoNet1* workflow.

*IsoNet2* also implements a mask-generation module that identifies empty regions in the tomogram where subtomograms should not be extracted. This improves training efficiency by avoiding unnecessary computation. In practice, mask generation could be performed on the denoised or CTF deconvolved tomograms using the default parameters before the Refine step.

### Processing of HIV dataset EMPIAR-10164

This dataset contains raw tilt series frames for immature HIV-1 dMACANC VLPs^15^. We chose 5 tomograms for testing. The raw movie frames were motion-corrected with *MotionCor2*^32^ and split into even and odd frames. The tilt series of even and odd images are stacked by “newstack” command in *IMOD*^33^. Full tilt-series alignment and defocus determination was performed with *AreTomo2*^34^ under default parameters. The alignment is then transferred to *IMOD* for tomogram reconstruction with weighted back projection algorithm. The tilt series are binned by 4 to reach the pixel size of 5.4 Å/pixel before tomogram reconstruction.

For *IsoNet2* processing, we first performed an initial *IsoNet2* Refine with missing-wedge weight of 200, cube size of 96 and network-based CTF correction with a B-factor of 0, using total 2,000 subtomograms or 400 subtomograms in each tomogram. The resulting network was then applied to the tomograms to generate the *IsoNet2*-processed tomograms. These tomograms were then subject to a mask generating process with the default parameter, *i.e.* std percentage of 80 and density percentage of 50 and z_crop of 0.2.

These masks are then used for a final *IsoNet2* Refine step with missing-wedge weight of 200, cube size of 128 and network-based CTF correction with a B-factor of 200 Å^2^, using total 5,000 subtomograms from 5 tomograms for 50 epochs. The final network model was then applied to the five tomograms. To estimate local resolution, a 160 pixel diameter subtomogram containing a spherical virus was cropped using the “crop volume” tool in *ChimeraX*^19^ and with contrast inverted by “vop multiply”. The local resolution was estimated for this cropped cubic subtomogram using *ResMap* v1.2^16^.

### Processing of 80S ribosome dataset EMPIAR-10985

This dataset contains cryoET data of 70S ribosomes imaged using *PACEtomo*^17^. Raw movie frames were motion-corrected with *MotionCor2* and split into even and odd frames. Tilt series of even and odd images were stacked by “newstack” command in *IMOD*. Full tilt-series alignment and defocus determination was performed with *AreTomo2* under default parameters, and the resulting alignment was imported into *IMOD* for tomogram reconstruction using the weighted back-projection algorithm. All tilt series were binned by 5 to reach a final pixel size of 5.35 Å/pixel prior to reconstruction.

For *IsoNet2* processing, we performed a refinement using the following parameters: missing-wedge weight = 200, cube size = 96, and network-based CTF correction with a B-factor of 300 Å^2^. A total of 2,400 subtomograms from 80 tomograms were used to train for 50 epochs. Local resolution was estimated by cropping a subtomogram (∼160 pixels in diameter) containing a well-centered ribosome particle using *ChimeraX*’s “crop volume” tool, followed by contrast inversion. Resolution estimation was performed with *ResMap* v1.2.

### Processing of FIB-milling cellular cryoET dataset EMPIAR-11830

This dataset contains tilt-series acquired from *Chlamydomonas reinhardtii* lamellae prepared by cryo-plasma FIB milling^18^. Defocus files and alignment files were obtained from the EMPIAR dataset. The alignment was imported into *IMOD* for tomogram reconstruction using the weighted back-projection algorithm. All tilt series were binned by 4× prior to reconstruction to reach a final pixel size of 7.84 Å/pixel.

For *IsoNet2* processing, 15 tomograms containing mitochondria were selected. We first conducted an *IsoNet2* Refine run using a missing-wedge weight of 20, cube size of 96, and CTF correction with a B-factor of 0. A total of 1,500 subtomograms (100 subtomograms per tomogram) were extracted and used for training. The resulting neural network was then applied to all 15 tomograms to generate the initial *IsoNet2*-processed volumes. Mask generation was performed using the default *IsoNet2* settings (standard-deviation percentile = 50%, density percentile = 50%, z-crop = 0.2).

These masks were used in a final *IsoNet2* Refine stage with missing-wedge weight of 200, cube size of 128, and B-factor of 0. The final refinement used 1,500 subtomograms for 100 epochs. The resulting trained network was then applied to 15 tomograms to yield the final *IsoNet2*-processed tomograms. To estimate local resolution, a subtomogram (128 pixels in diameter) was cropped using *ChimeraX*’s “crop volume” tool. The contrast was inverted for this volume in *ChimeraX*, and the localized resolution was estimated using *ResMap* v1.2.

### Processing of 80S ribosome dataset EMPIAR-10045

This dataset consists of tilt-series of mammalian 80S ribosomes^35^. The tilt series were split into even and odd based on the tilt angles. Defocus values were provided alongside the dataset. *IMOD* was used for tomogram reconstruction using weighted back-projection. All seven tilt series were binned by 6× to obtain a final pixel size of 13.02 Å/pixel.

*IsoNet2* processing refinement using missing-wedge weight of 20, cube size of 96, and network-based CTF correction with B-factor of 0, using a training set of 3,000 subtomograms (500 subtomograms per tomogram) for 50 epochs. The final network was applied to the tomograms to obtain the final *IsoNet2*-processed volumes. The *DeepDeWedge*-processed tomogram was downloaded from the data accompanying their paper.

### Processing cellular cryoET dataset EMPIAR-11078

This dataset contains tilt-series of *in situ* cryoET of the *Chlamydomonas reinhardtii* ciliary transition zone^36^. Raw movie frames were motion-corrected with *MotionCor2* and split into even and odd frame stacks. Defocus values were obtained from the information from EMPIAR dataset. Tilt-series alignment was performed in *AreTomo2* using default settings, and alignments were imported into *IMOD* for tomogram reconstruction using the weighted back-projection method. All tilt series were binned by 3× to achieve a final pixel size of 10.26 Å/pixel.

*IsoNet2* initial processing was performed on 8 selected tomograms with a missing-wedge weight of 20, cube size of 96, and B-factor of 0, trained on 2,400 subtomograms (300 subtomograms per tomogram). The resulting model was applied to the tomograms to generate the initial *IsoNet2*-processed volumes. Mask generation was conducted using *IsoNet2*’s default settings (standard-deviation percentile = 50%, density percentile = 50%, z-crop = 0.2).

A final *IsoNet2* Refine was then performed with missing-wedge weight of 200, box size of 128, and B-factor of 0, using 2,400 subtomograms and 50 training epochs. The final network was applied to all tomograms. To compare with the result from *DeepDeWedge*, the *DeepDeWedge*-processed tomogram was downloaded from the data accompanied with their paper.

### Tomogram segmentation and rendering

Segmentation of ribosomes was carried out by fitting the *Escherichia coli* 70S ribosome atomic model^18^ (PDB: 7K00) into each corresponding density in the cryoET tomograms using the “fit in map” tool in *ChimeraX*. Immersive inspection with the AR/VR visualization tools facilitated precise assessment of model placement in 3D. After fitting, each ribosome was colored using the “Color Zone” tool to delineate individual particles and highlight subunit-specific features.

Segmentations for the HIV and cellular cryoET datasets shown in Figures 1 and 3 were performed manually in *Dragonfly* (Comet Group). Viral particles, microtubules, and protein components were segmented using the Otsu brush in *Dragonfly*, whereas membranes were traced using the line-segment tool. The resulting segmentations were exported as TIFF volumes and imported into *ChimeraX*.

The raw densities rather than segmented volumes are displayed in our rendering. To define regions for raw densities, segmentation volumes were low-pass filtered (Gaussian, σ=20 voxels) to generate smooth masks, then binarized in *ChimeraX*. Masks were applied using “vop multiply” to isolate individual components from the original tomograms while preserving complementary density.

For the cellular tomogram in Figure 3, all density was displayed using a single global threshold, with distinct colors assigned to structural classes to produce a dense, Goodsell-style visualization. The ATP synthase atomic model^22^ (PDB: 6RD4) was fitted directly into the *IsoNet2*-processed tomograms, and proteasomes were interpreted using the 26S model^23^ (PDB: 4CR2).

### Data availability

The five datasets used in the paper are publicly available through: EMPIAR: EMPIAR-10164 for HIV particles in Fig.1; EMPIAR-10985 for 70S ribosome in Fig.2; EMPIAR-11830 for Chlamydomonas reinhardtii lamellae prepared by cryo-plasma FIB milling in Fig.3; EMPIAR-11078 and EMPIAR 10045 are cellular and ribosome dataset for Extended Data Fig.3.

### Code availability

The code is available at https://github.com/IsoNet-cryoET/IsoNet2 with both code and tutorial.

## Acknowledgements

We thank Alex Wu for testing the software. This research has been supported by the National Institute of General Medicine of the National Institutes of Health (R01GM071940 to Z.H.Z.).

## Author information

### Contributions

Y.T.L designed the algorithm, implemented the software, performed the experiments, prepared the figures, and wrote the manuscript. H.F. contributed to method development, processed the data, interpreted the results, and helped writing documentation. J.J. prepared the figures, contributed to the code, and wrote the manuscript. L.T. contributed to coding and data processing and made documentation.

X.Z. designed and implemented the GUI. Z.H.Z. supervised and managed the project.

### Ethics declarations Competing interests

The authors declare no competing interests.

### Additional information

Extended data is available for this paper.

---

**Extended Data Fig. 1.**
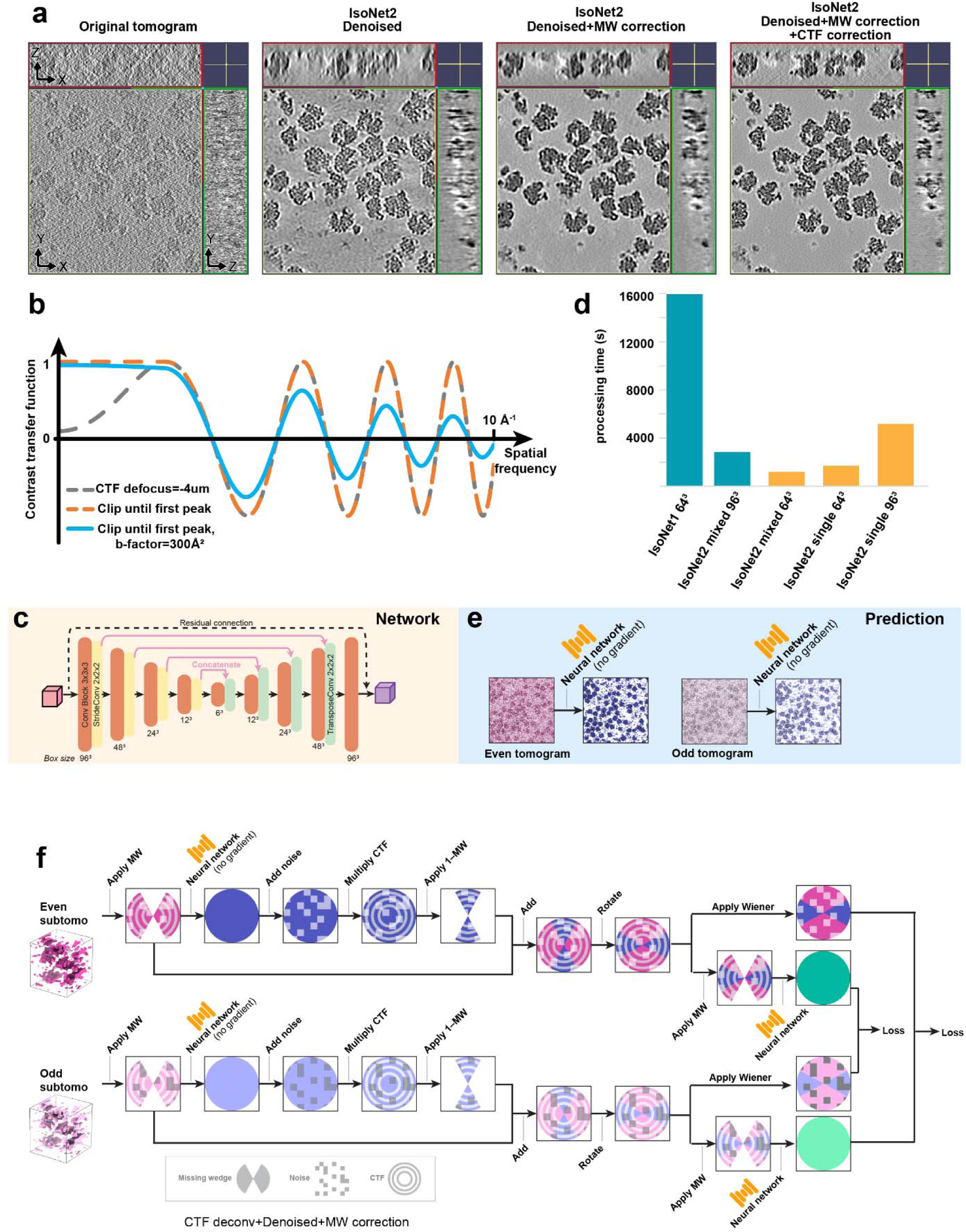
Overview of *IsoNet2* processing. **a,** Sequential improvements in tomogram quality for the ribosome tomograms produced by *IsoNet2*. From left to right: original tomogram, denoised result, denoised + missing-wedge–corrected output, and full denoising + missing-wedge + CTF–corrected reconstruction. **b,** *IsoNet2*’s CTF curve. Low-frequency CTF components are clipped to the first peak, preventing incorrect modulation at very low resolution. Optional B-factor weighted CTF is in blue. **c,** *IsoNet2* network architecture: a 3D U-Net with four down- sampling and up-sampling layers, residual connections, and a larger receptive field (96³ input cube). **d,** Processing time comparison between *IsoNet1* and *IsoNet2* under different training modes and cube sizes. Mixed-precision *IsoNet2* achieves ∼10× faster computation than *IsoNet1* and enables efficient training on larger boxes. **e,** Prediction pipeline in which even and odd tomograms are independently processed by the trained neural network to generate refined volumes. **f,** *IsoNet2* Refine step but with CTF correction using Wiener filter.

**Extended Data Fig. 2.**
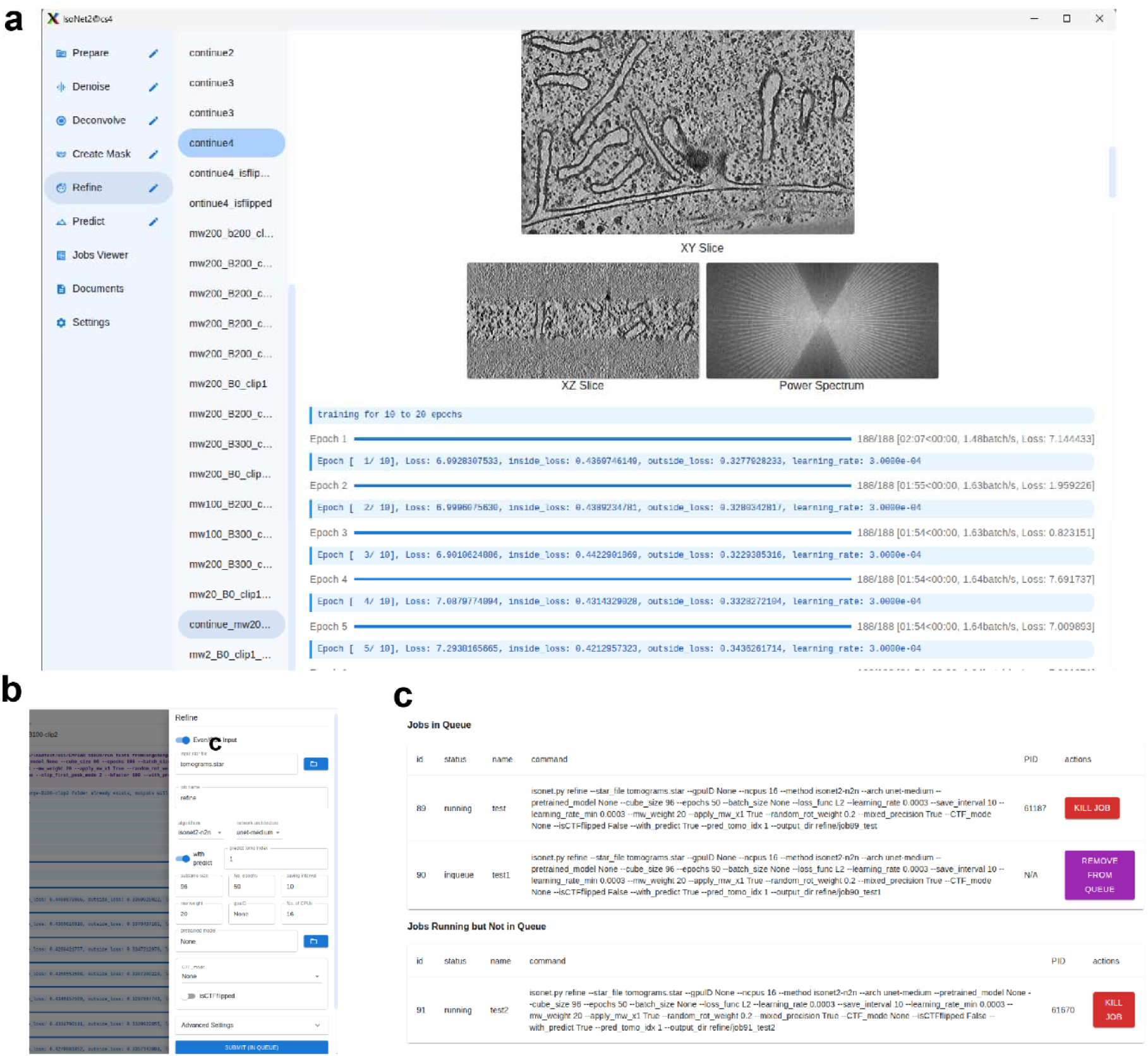
*IsoNet2* graphical user interface and job-management system. **a,** The *IsoNet2* desktop interface, built on modern web-based frameworks, provides an integrated environment for tomogram processing. **b,** The job submission panel. **c,** The *IsoNet2* job manager.

**Extended Data Fig. 3.**
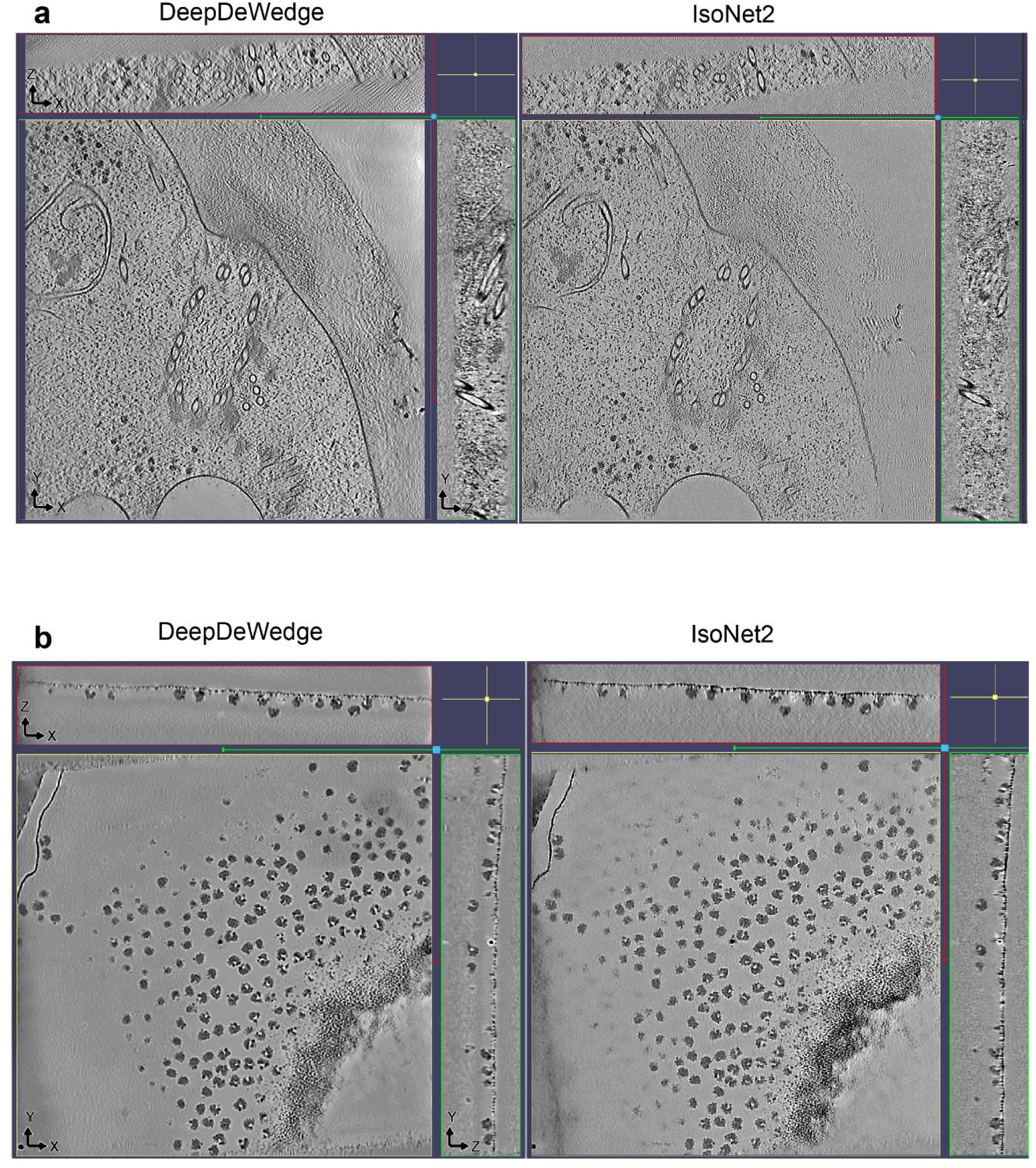
Comparison of *IsoNet2* and *DeepDeWedge*-processed tomograms. Orthogonal slices of a tomogram in EMPIAR-11078 (upper panel) and a tomogram in EMPIAR-10045 (lower panel) processed with *DeepDeWedge* and *IsoNet2*.

**Extended Data Fig. 4.**
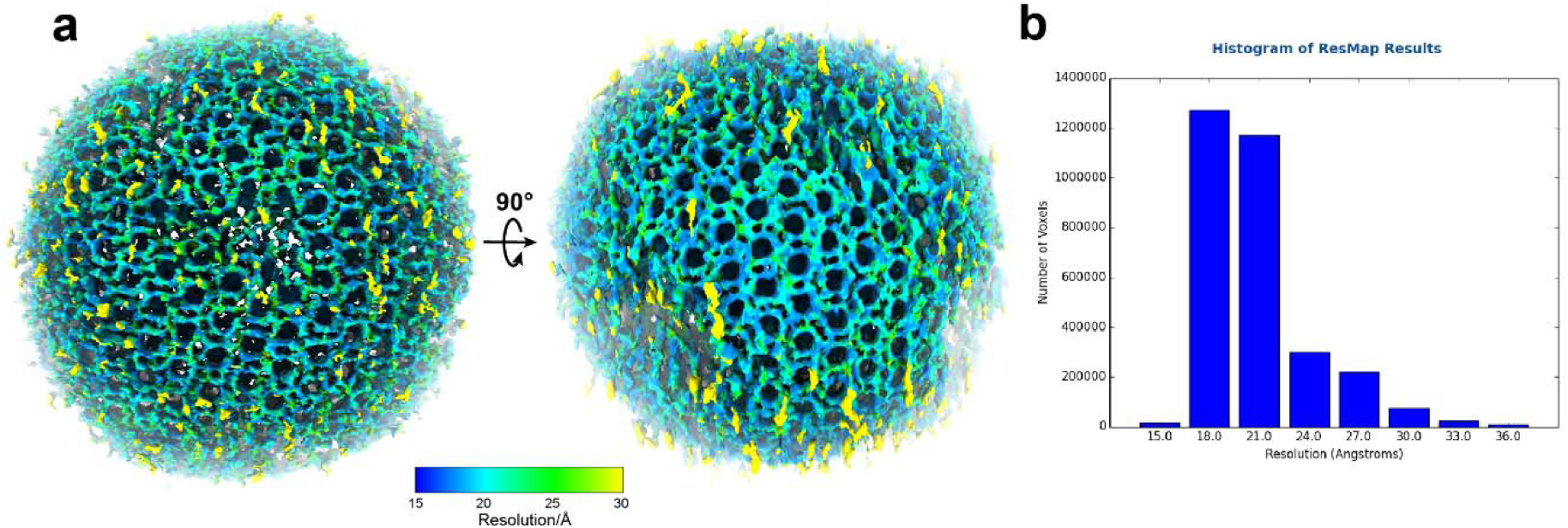
Local resolution estimation for an HIV particle **a,** Surface rendering of an HIV particle from EMPIAR-10164 by its local resolution. **b,** Histogram showing the number of pixels across the map resolution range.

**Extended Data Fig. 5.**
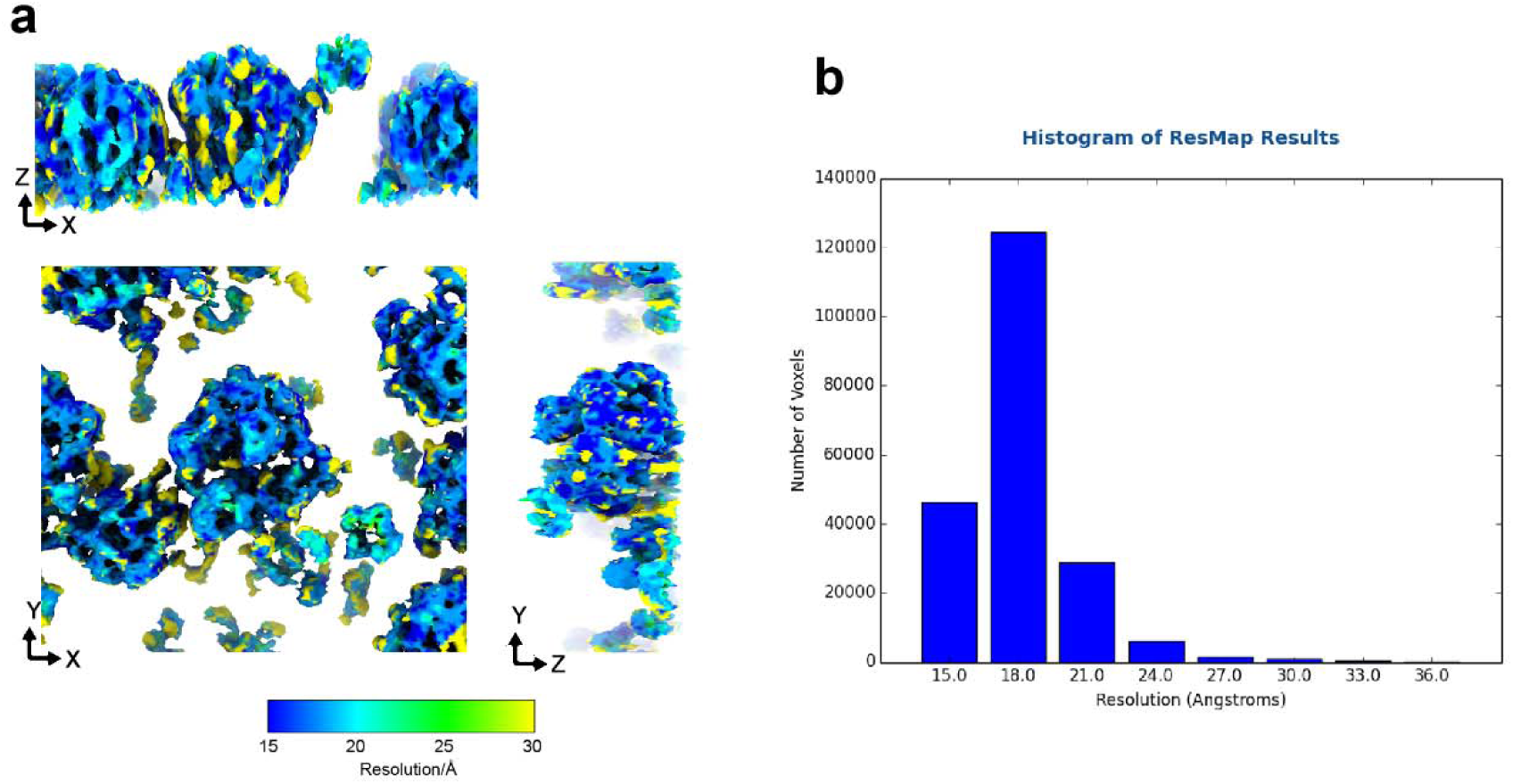
Local resolution estimation for 80S ribosomes **a,** Surface rendering of 80S ribosomes from EMPIAR-10985 by local resolution. **b,** Histogram showing the number of pixels across the map resolution range.

**Extended Data Fig. 6.**
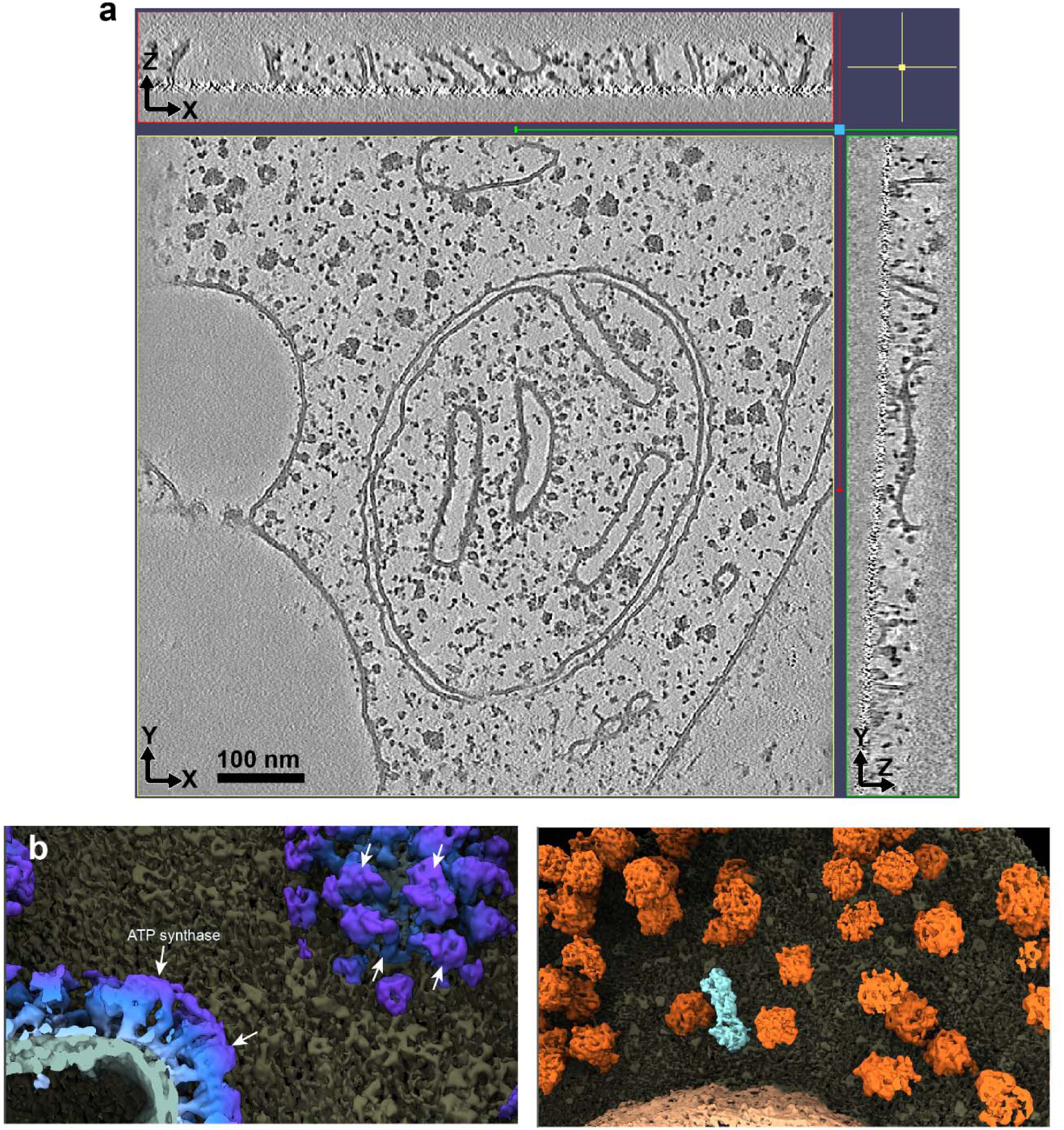
*IsoNet2* corrected cellular tomogram **a-b**, Orthogonal slices (**a)** for the tomogram shown in Fig. 3 with more 3D details of the rendering shown in **b.**

**Extended Data Fig. 7.**
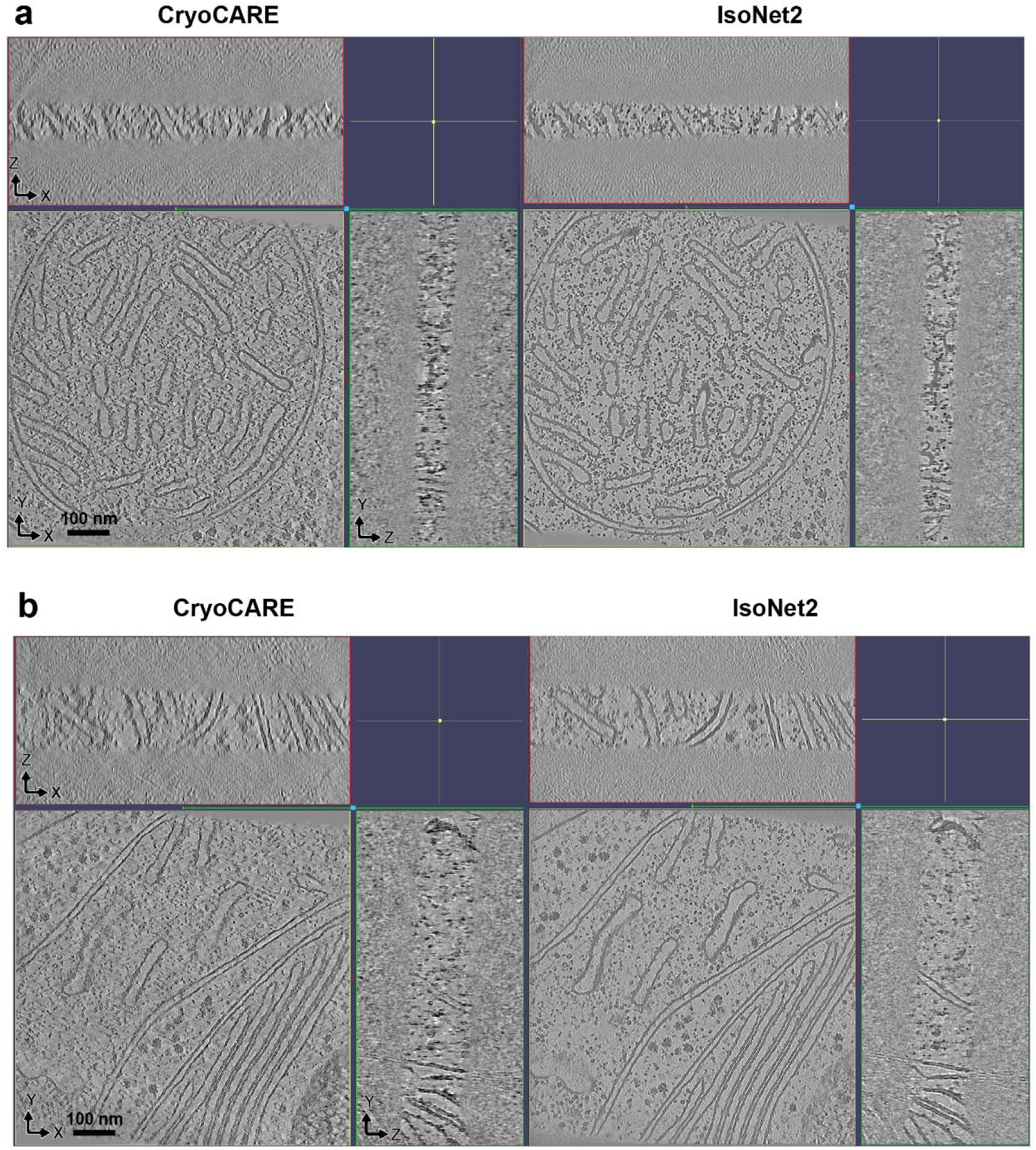
Comparison of *IsoNet2*- and *CryoCARE*-processed tomograms Orthogonal slices of tomograms in EMPIAR-10985 processed with *CryoCARE* and *IsoNet2*

**Extended Data Fig. 8.**
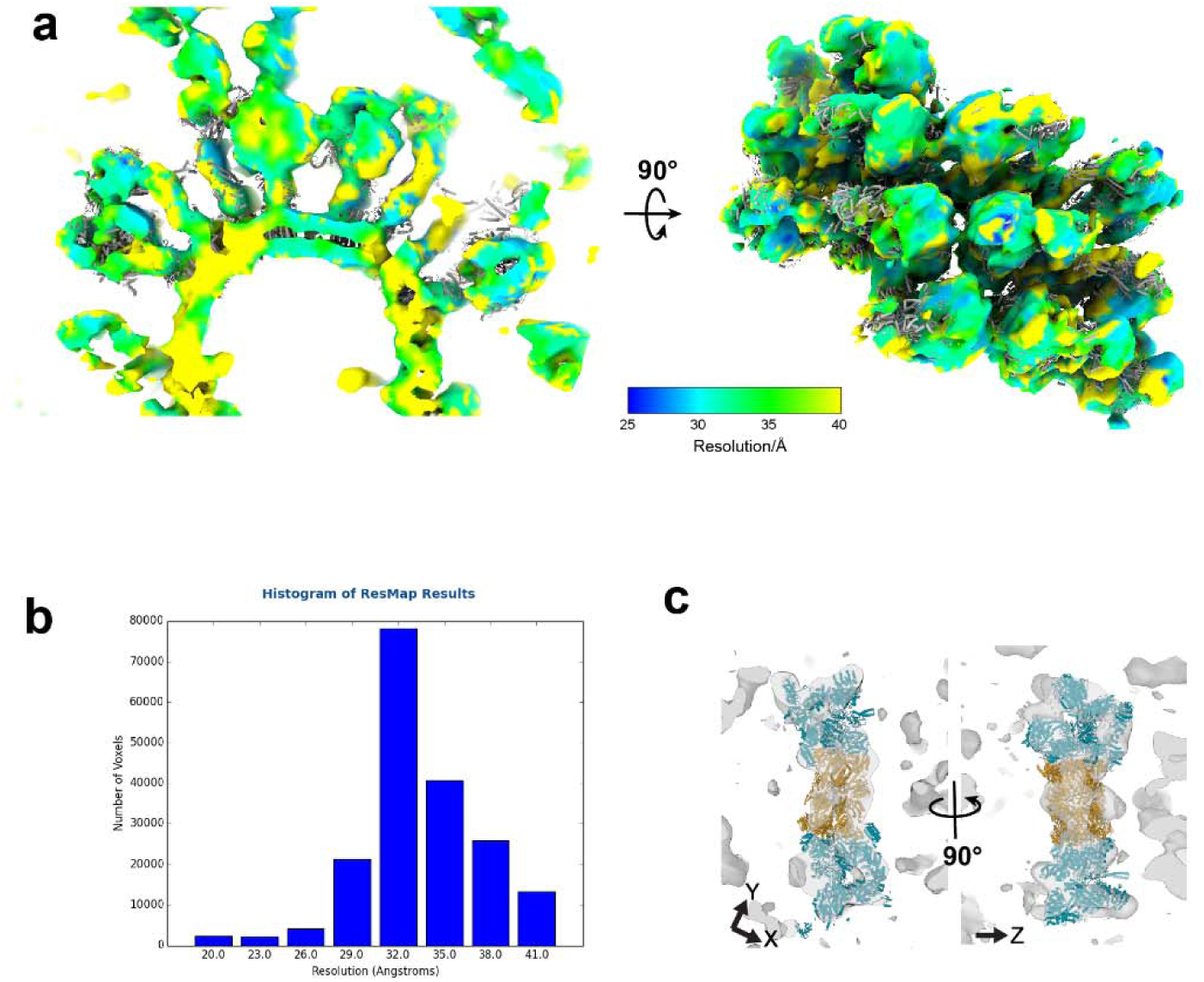
Local resolution estimation for FIB-milled cellular tomogram **a**, Surface rendering of the ATP synthase super-complexes from EMPIAR-11830 by local resolution. **b**, Histogram showing the number of pixels across the map resolution range. **c**, Proteosome density (different from Fig. 3D) showing atomic model fitting into the cryoET density in transparent gray.

## Notes

### Competing Interest Statement

The authors have declared no competing interest.

## References

1. Gilbert, P. Iterative methods for the three-dimensional reconstruction of an object from projections. J. Theor. Biol. 36, 105–117 (1972).

2. Andersen, A. H. & Kak, A. C. Simultaneous algebraic reconstruction technique (SART): a superior implementation of the art algorithm. Ultrason. Imaging 6, 81–94 (1984).

3. Buchholz, T.-O. et al. Content-aware image restoration for electron microscopy. Methods Cell Biol. 152, 277–289 (2019).

4. Lehtinen, J., et al. Noise2Noise: Learning Image Restoration without Clean Data. Preprint at 10.48550/arXiv.1803.04189 (2018).

5. Liu, Y.-T. et al. Isotropic reconstruction for electron tomography with deep learning. Nat. Commun. 13, 6482 (2022).

6. Wiedemann, S. & Heckel, R. A deep learning method for simultaneous denoising and missing wedge reconstruction in cryogenic electron tomography. Nat. Commun. 15, 8255 (2024).

7. Kishore, V., et al. Efficient reconstruction and denoising of cryo-ET data with end-to-end localized deep learning. Preprint at 10.48550/arXiv.2501.15246 (2025).

8. Rather, I. H., Kumar, S. & Gandomi, A. H. Breaking the data barrier: a review of deep learning techniques for democratizing AI with small datasets. Artif. Intell. Rev. 57, 226 (2024).

9. Unnisa, Z. et al. Impact of fine-tuning parameters of convolutional neural network for skin cancer detection. Sci. Rep. 15, 14779 (2025).

10. Tegunov, D., Xue, L., Dienemann, C., Cramer, P. & Mahamid, J. Multi-particle cryo-EM refinement with M visualizes ribosome-antibiotic complex at 3.5 Å in cells. Nat. Methods 18, 186–193 (2021).

11. Bepler, T., Kelley, K., Noble, A. J. & Berger, B. Topaz-Denoise: general deep denoising models for cryoEM and cryoET. Nat. Commun. 11, 5208 (2020).

12. Scheres, S. H. W. RELION: Implementation of a Bayesian approach to cryo-EM structure determination. J. Struct. Biol. 180, 519–530 (2012).

13. Micikevicius, P., et al. Mixed Precision Training. Preprint at 10.48550/arXiv.1710.03740 (2018).

14. Ronneberger, O., Fischer, P. & Brox, T. U-Net: Convolutional Networks for Biomedical Image Segmentation. Preprint at 10.48550/arXiv.1505.04597 (2015).

15. Schur, F. K. M. et al. An atomic model of HIV-1 capsid-SP1 reveals structures regulating assembly and maturation. Science 353, 506–508 (2016).

16. Kucukelbir, A., Sigworth, F. J. & Tagare, H. D. Quantifying the local resolution of cryo- EM density maps. Nat. Methods 11, 63–65 (2014).

17. Eisenstein, F. et al. Parallel cryo electron tomography on in situ lamellae. Nat. Methods 20, 131–138 (2023).

18. Watson, Z. L. et al. Structure of the bacterial ribosome at 2 Å resolution. eLife 9, e60482 (2020).

19. Goddard, T. D. et al. UCSF ChimeraX: Meeting modern challenges in visualization and analysis. Protein Sci. Publ. Protein Soc. 27, 14–25 (2018).

20. Kelley, R. et al. Towards community-driven visual proteomics with large-scale cryo- electron tomography of Chlamydomonas reinhardtii. 2024.12.28.630444 Preprint at 10.1101/2024.12.28.630444 (2024).

21. Goodsell, D. S. & Lasker, K. Integrative visualization of the molecular structure of a cellular microdomain. Protein Sci. Publ. Protein Soc. 32, e4577 (2023).

22. Dietrich, L., Agip, A.-N. A., Kunz, C., Schwarz, A. & Kühlbrandt, W. In situ structure and rotary states of mitochondrial ATP synthase in whole Polytomella cells. Science 385, 1086–1090 (2024).

23. Unverdorben, P. et al. Deep classification of a large cryo-EM dataset defines the conformational landscape of the 26S proteasome. Proc. Natl. Acad. Sci. 111, 5544–5549 (2014).

24. Abramson, J. et al. Accurate structure prediction of biomolecular interactions with AlphaFold 3. Nature 630, 493–500 (2024).

25. Beck, M. & Baumeister, W. Cryo-Electron Tomography: Can it Reveal the Molecular Sociology of Cells in Atomic Detail? Trends Cell Biol. 26, 825–837 (2016).

26. Paszke, A., et al. Automatic differentiation in PyTorch. in (2017).

27. Abadi, M. et al. TensorFlow: a system for large-scale machine learning. in *Proceedings of the 12th USENIX conference on Operating Systems Design and Implementation* 265–283 (USENIX Association, USA, 2016).

28. Burt, A. et al. An image processing pipeline for electron cryo-tomography in RELION-5. 2024.04.26.591129 Preprint at 10.1101/2024.04.26.591129 (2024).

29. Liu, Y.-T., Fan, H., Hu, J. J. & Zhou, Z. H. Overcoming the preferred-orientation problem in cryo-EM with self-supervised deep learning. Nat. Methods 22, 113–123 (2025).

30. Zhang, K. Gctf: Real-time CTF determination and correction. J. Struct. Biol. 193, 1–12 (2016).

31. Tegunov, D. & Cramer, P. Real-time cryo-electron microscopy data preprocessing with Warp. Nat. Methods 16, 1146–1152 (2019).

32. Zheng, S. Q. et al. MotionCor2: anisotropic correction of beam-induced motion for improved cryo-electron microscopy. Nat. Methods 14, 331–332 (2017).

33. Mastronarde, D. N. & Held, S. R. Automated tilt series alignment and tomographic reconstruction in IMOD. J. Struct. Biol. 197, 102–113 (2017).

34. Zheng, S. et al. AreTomo: An integrated software package for automated marker-free, motion-corrected cryo-electron tomographic alignment and reconstruction. J. Struct. Biol. X 6, 100068 (2022).

35. Bharat, T. A. M. & Scheres, S. H. W. Resolving macromolecular structures from electron cryo-tomography data using subtomogram averaging in RELION. Nat. Protoc. 11, 2054–2065 (2016).

36. van den Hoek, H., et al. In situ architecture of the ciliary base reveals the stepwise assembly of intraflagellar transport trains. Science 377, 543–548 (2022).

